# Differential regulation of native and learned behavior by creb-1/crh-1 in Caenorhabditis elegans

**DOI:** 10.1101/508986

**Authors:** Yogesh Dahiya, Saloni Rose, Shruti Thapliyal, Shivam Bhardwaj, Maruthi Prasad, Kavita Babu

## Abstract

Memory formation is crucial for the survival of animals. Here, we study the effect of different ***crh-1*** (***C. elegans*** homolog of mammalian CREB1) mutations on the ability of ***C. elegans*** to form long-term memory (LTM). Null mutants in ***creb1/crh-1*** are defective in LTM formation across phyla. We show that specific isoforms of CREB1/CRH-1, CRH-1c and CRH-1e, are primarily responsible for memory related functions of the transcription factor in ***C. elegans***. Silencing of CRH-1e expressing neurons during training for LTM formation abolishes the long-term memory of the animal. Further, CRH-1e expression in RIM or AVE neurons is sufficient to rescue long-term memory defects of ***creb1/crh-1*** null mutants. We show that apart from being LTM defective, ***creb1/crh-1*** null mutant animals show defects in native chemotaxis behavior. We characterize the amino acids K247 and K266 as responsible for the LTM related functions of CRH-1 while being dispensable for it’s native chemotaxis behavior. These findings provide insight into the spatial and temporal workings of a crucial transcription factor and can be further exploited to find CREB1 targets involved in the process of memory formation.

## 2. Introduction

Native behaviors are important to animals and many of these behaviors can be modified through experience. Native behaviors, in part, represent the hardwired neural circuitry put in place during the development of an animal (Tierney, 1986; Bateson and Mameli, 2007). Mutations that affect the native behavior could also alter the ability of animals to modify that particular behavior in response to experience. Hence, learning deficits that are attributed to certain alleles could sometimes arise due to underlying native behavioral defects. Here, we show the effect of different ***crh-1*** (homolog of mammalian CREB1) mutations on native and learned chemotaxis behaviors in ***Caenorhabditis elegans.***

***C. elegans*** are soil dwelling nematodes that live on microbes as their food source. Their nervous system consists of 302 neurons. They show a variety of robust and adaptable behaviors like chemotaxis and thermotaxis (Hart, 2006; Ardiel and Rankin, 2010). ***C. elegans*** move in a sinusoidal wave pattern. Under laboratory conditions, their continuous forward movement is punctuated by frequent stops and events of backward movement called as reversals. After each reversal, the probability of change in direction of movement increases significantly. One of the strategies deployed by ***C. elegans*** to move in response to a chemical gradient is to change the frequency of reversals in response to the gradient (Pierce-Shimomura et al., 1999).

CREB1 (cAMP Response Element binding Protein 1), was initially characterized for its role in long-term facilitation of the gill withdrawal reflex in ***Aplysia*** (Dash et al., 1990). Neural activity dependent CREB1 activation in a defined set of neurons is one of the requirements for the process of long-term memory (LTM) formation (Flavell and Greenberg, 2008). CREB1 null mutants have been shown to be defective in LTM formation across phyla (Silva et al., 1998).

***C. elegans*** have native preferences for certain type of odors. They detect metabolic byproducts of microbes as cues for finding their food. For example, under well fed naïve condition worms are attracted to AWC sensed odorants like isoamyl alcohol (Bargmann, 2006). However, this odor to behavior relationship can be modified if worms are exposed to periods of starvation along-with chemo-attractants (Pereira and Kooy, 2012). Likewise, neutral concentrations of butanone can be turned attractive when repetitively paired with food. These modified behaviors persist for 40 hours after training (Kauffman et al., 2011).

The role of ***crh-1*** in long-term memory formation has been described utilizing variety of training paradigms using the ***crh-1*** null mutant strain ***crh-1(tz2)*** (Kauffman et al., 2010; Nishida et al., 2011; Timbers and Rankin, 2011; Lau et al., 2013; Xin et al., 2016). Chemotaxis behavior is generally used to assess the performance of memory formation and retrieval assays. Chemotaxis in ***C. elegans*** is dependent on a large number of subtle behaviors like reversals and head bending (Pierce-Shimomura et al., 1999; Iino and Yoshida, 2009; Larsch et al., 2015). We have observed that the innate ability of worms to modulate reversals in response to external stimuli is a critical factor in determining the performance of ***C. elegans*** in chemotaxis assays, thereby affecting the readout of learning assays.

The role of ***crh-1*** in defining long-term memory in ***C. elegans*** has only been partially understood. First, given its pleiotropy of function, it is unclear which ***crh-1*** isoforms are involved in memory related functions. Second, the motifs on CRH-1 protein specifically involved in memory related processes are unknown. Third, the identity of neurons where ***crh-1*** is required for memory formation is largely unknown. Here, we used CRISPR based mutagenesis to dissect out memory related functions of CREB1/CRH-1. Our experiments show that two of the six potential isoforms of CRH-1, CRH-1c and CRH-1e, are responsible for these functions. Further, amino acid residues Lys247 and Lys266 are critical for memory related functions of CRH-1 while being dispensable for native chemotaxis behavior. We have also identified AVE and/or RIM as the minimal set of neurons required for memory related functions of CRH-1.

## 3. Materials and Methods

### 3.1 Strains

***C. elegans*** were maintained using standard methods (Brenner, 1974). Wild-type N2 and ***crh-1(tz2)*** lines were obtained from the ***Caenorhabditis*** Genetics Center (University of Minnesota, Minneapolis, MN). Strains used in this study are listed in 3.

### 3.2 Constructs and Transgenes

The ***crh-1*** isoforms (***crh-1a-f***) were cloned into the pPD49.26 vector backbone. ***crh-1b*** and ***crh-1c*** cDNAs were synthesized from Sigma-Aldrich and the rest were obtained by reverse transcription and PCR using wild type ***C. elegans*** RNA. The N2 (WT) or ***crh-1(tz2)*** mutant lines were used for transforming the CRH-1 isoforms. Transformations were done as described previously (Mello et al., 1991). The rescue constructs and the promoter fusion constructs were injected in concentrations of 10-20ng/µl. p***myo-2***::mCherry (2ng/µl) or p***vha-6***::mCherry (10ng/µl) or p***unc-122::***GFP (25ng/µl) were used as co-injection markers. mCherry and GFP cDNA were cloned from pCFJ90 and pPD95.75 respectively. The primers and plasmids used for the different cloning experiments are tabulated in Table 1 and 2 respectively.

**Table 1:**
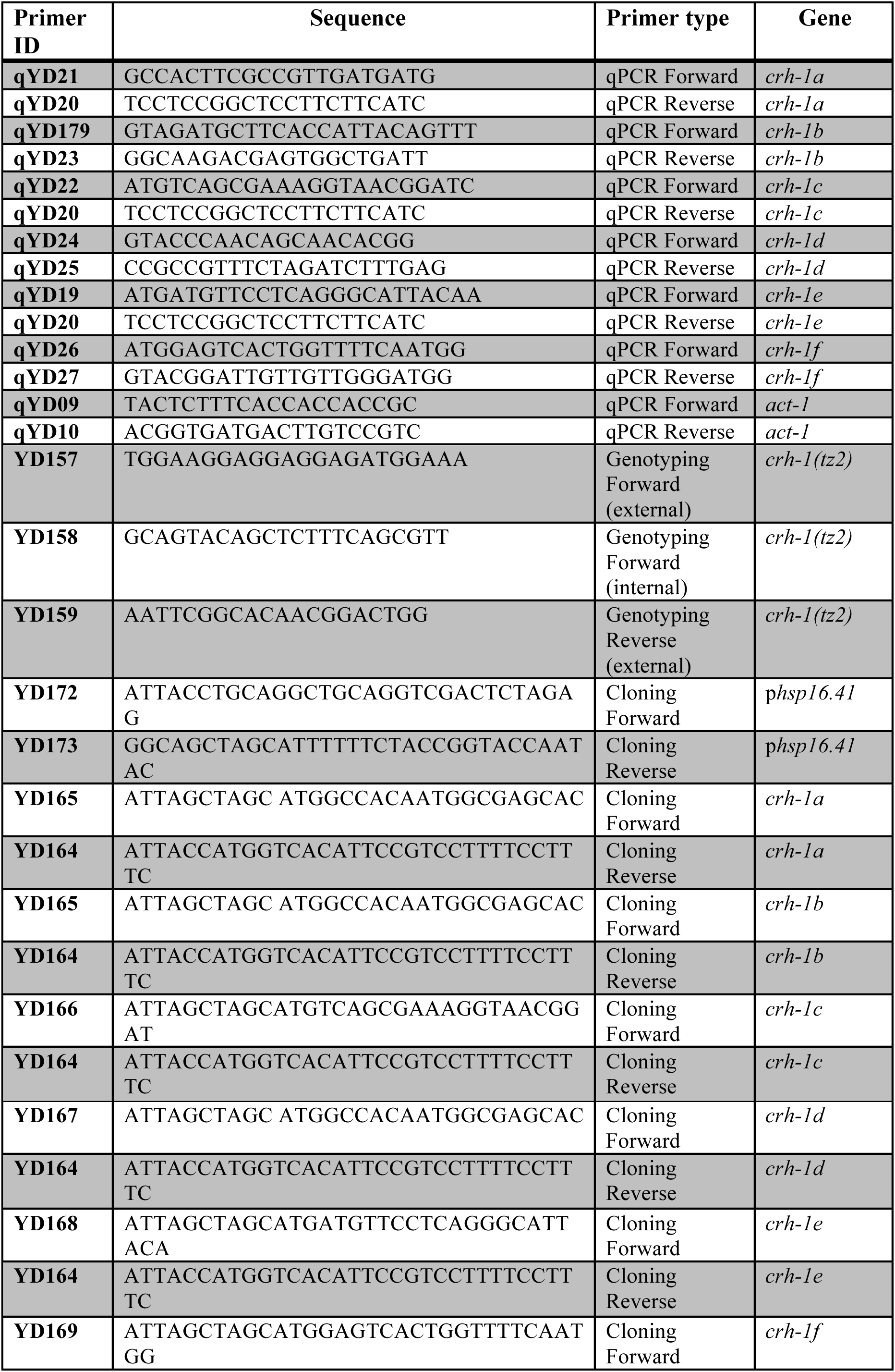
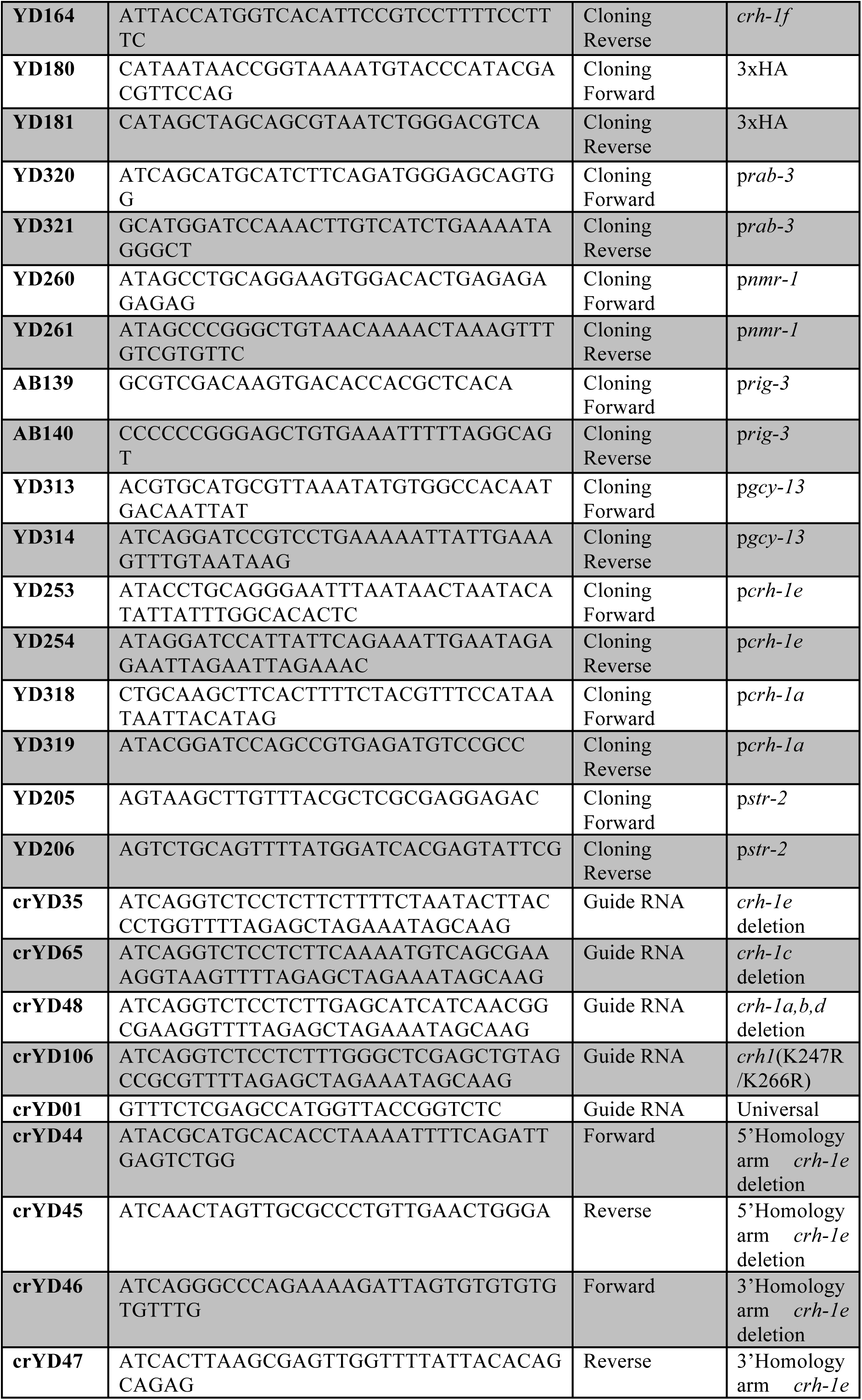
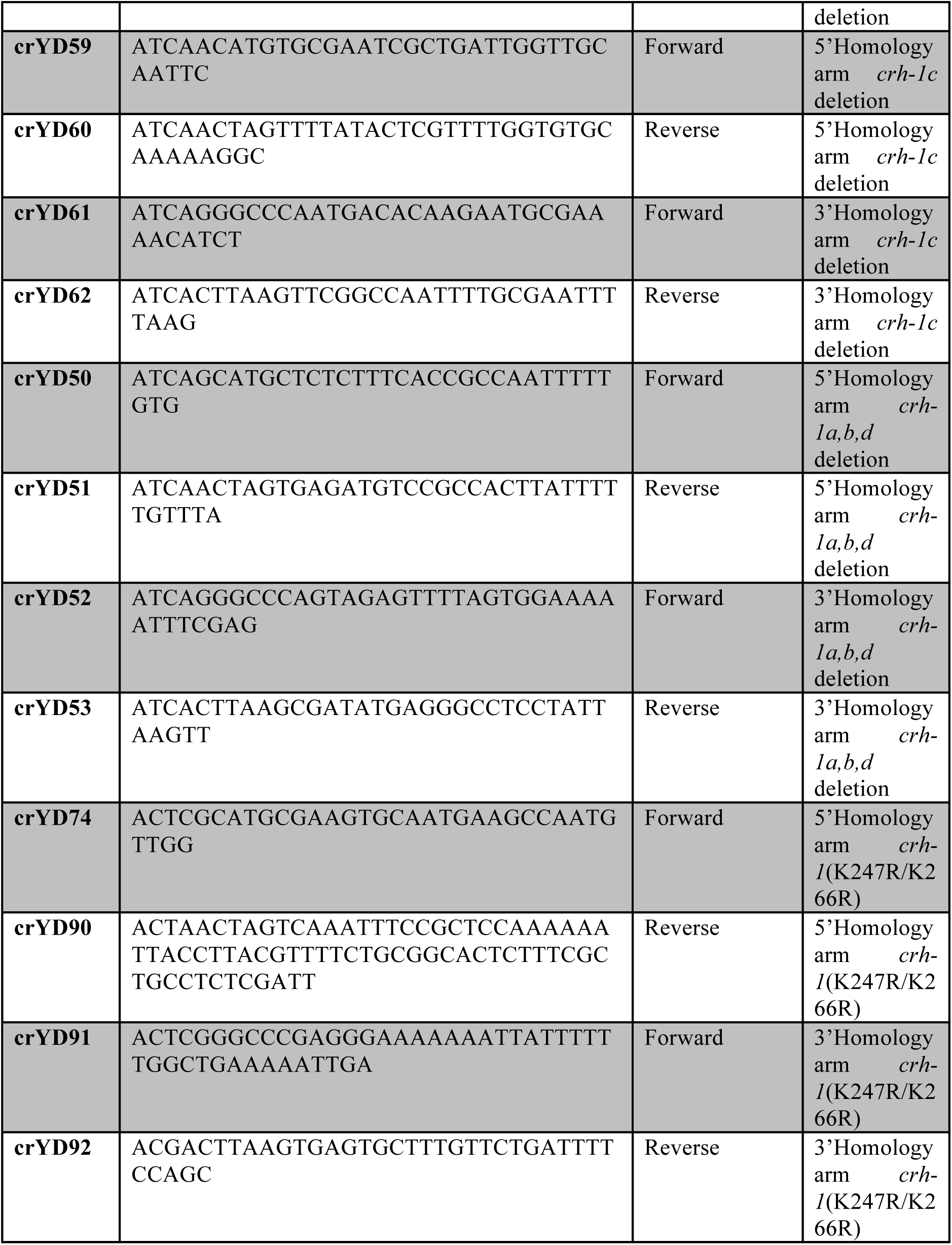
List of primers used in this study

**Table 2:** List of plasmids used in this study

| S. no. | Plasmid ID | Plasmid |
| --- | --- | --- |
| 1 | pBAB713 | *prab-3::CRH-1a* |
| 2 | pBAB714 | *prab-3::CRH-1b* |
| 3 | pBAB715 | *prab-3::CRH-1c* |
| 4 | pBAB716 | *prab-3::CRH-1d* |
| 5 | pBAB717 | *prab-3::CRH-1e* |
| 6 | pBAB719 | *prab-3::CRH-1f* |
| 7 | pBAB712 | *phsp16.41::CRH-1e* |
| 8 | pBAB726 | *pnmr-1::CRH-1e* |
| 9 | pBAB746 | *pcrh-1e::CRH-1e* |
| 10 | pBAB748 | *prig-3::mCherry* |
| 11 | pBAB749 | *popt-3::mCherry* |
| 12 | pBAB750 | *pgcy-13::mCherry* |
| 13 | pBAB747 | *prig-3::CRH-1e* |
| 14 | pBAB745 | *popt-3::CRH-1e* |
| 15 | pBAB744 | *pgcy-13::CRH-1e* |
| 16 | pBAB761 | *pstr-2::ChR2::YFP* |
| 17 | pBAB751 | *pcrh-1e::GFP* |
| 18 | pBAB736 | *pnmr-1::mCherry* |
| 19 | pBAB752 | *pcrh-1a::GFP* |
| 20 | pBAB741 | *pcrh-1e::HisC11::sl2::GFP* |
| 21 | pBAB709 | Repair template *crh-1a,b,d* deletion |
| 22 | pBAB711 | Repair template *crh-1c* deletion |
| 23 | pBAB708 | Repair template *crh-1e* deletion |
| 24 | pBAB760 | Repair template *crh-1K247R/K266R* point mutation |
| 25 | pBAB757 | Repair template 3xHA::CRH-1e |
| 26 | pBAB758 | Repair template 3xHA::CRH-1c |
| 27 | pBAB759 | gRNA *crh-1a,b,d* deletion |
| 28 | pBAB762 | gRNA *crh-1c* deletion |
| 29 | pBAB763 | gRNA *crh-1e* deletion |
| 30 | pBAB756 | gRNA *crh-1K247R/266R* point mutation |

**Table 3:** List of Strains used in this study

| Strain | Genotype | Description |
| --- | --- | --- |
| BAB701 | *crh-1(tz2)* (CGC strain YT17) | From CGC (outcrossed 2X) |
| BAB713 | *crh-1; prab-3::CRH-1a* | Array no. IndEx713 |
| BAB714 | *crh-1; prab-3::CRH-1b* | Array no. IndEx714 |
| BAB715 | *crh-1; prab-3::CRH-1c* | Array no. IndEx715 |
| BAB716 | *crh-1; prab-3::CRH-1d* | Array no. IndEx716 |
| BAB717 | *crh-1; prab-3::CRH-1e* | Array no. IndEx717 |
| BAB708 | CRISPR based deletion corresponding to the first exon of *crh-1e* mRNA | *crh-1*(Ind708+loxP) |
| BAB709 | CRISPR deletion corresponding to the first exon of *crh-1a,b,d* mRNAs | *crh-1*(Ind709+loxP) |
| BAB710 | CRISPR based deletion corresponding to the first exon of *crh-1c* and *crh-1e* mRNAs | *crh-1*(Ind710+loxP) |
| BAB711 | CRISPR based deletion corresponding to the first exon of *crh-1c* mRNA | *crh-1*(Ind711+loxP) |
| BAB719 | *prab-3::CRH-1f* | Integrated line IndIs719 |
| BAB726 | *crh-1; pnmr-1::CRH-1e* | Array no. IndEx726 |
| BAB736 | *pnmr-1::mCherry; pcrh-1e::GFP* | Array no. IndEx736 |
| BAB703 | *prig-3::mCherry; pcrh-1e::GFP* | Array no. IndEx703 |
| BAB704 | *popt-3::mCherry; pcrh-1e::GFP* | Array no. IndEx704 |
| BAB705 | *pgcy-13::mCherry; pcrh-1e::GFP* | Array no. IndEx705 |
| BAB747 | *crh-1; prig-3::CRH-1e* | Array no. IndEx747 |
| BAB745 | *crh-1; popt-3::CRH-1e* | Array no. IndEx745 |
| BAB744 | *crh-1; pgcy-13::CRH-1e* | Array no. IndEx744 |
| BAB746 | *crh-1; pcrh-1e::CRH-1e* | Array no. IndEx746 |
| BAB741 | *pcrh-1e::HISC11::SL2::GFP* | Array no. IndEx741 |
| BAB712 | *crh-1; phsp16.41::CRH-1e* | Array no. IndEx712 |
| BAB754 | CRISPR based addition of 3xHA::CRH-1c and e | *crh-1*(Ind754 [3xHA::*crh-1c*+loxP; 3xHA::*crh-1e*+loxP]) |
| BAB760 | CRISPR based substitution of K247R/266R | *crh-1*(Ind760 [K247R/K266R]+loxP) |
| BAB735 | *pnmr-1::mCherry; pcrh-1a::GFP* | Array no. IndEx735 |
| BAB761 | *pstr-2::ChR2::YFP; prig-3::GCaMP5G* | Array no. IndEx761 |
| BAB762 | BAB760; *pstr-2::ChR2::YFP; prig-3::GCaMP5G* | *crh-1*(Ind760 [K247R/K266R]+loxP) with array |

### 3.3 Behavioral assays

#### Conditioning with isoamyl alcohol (IAA) and high temperature

A dry heating block was maintained at 37°C. 20µl of 10% IAA (diluted in ethanol) was kept on a piece of cover glass on the heating block. A petri-plate containing ***C. elegans*** on ***E.Coli*** OP50 lawn was inverted onto the setup for 2 minutes (m). The training cycle was repeated 5 times with an inter-training interval of 10m (illustrated in Figure 1A).

**Figure 1.**
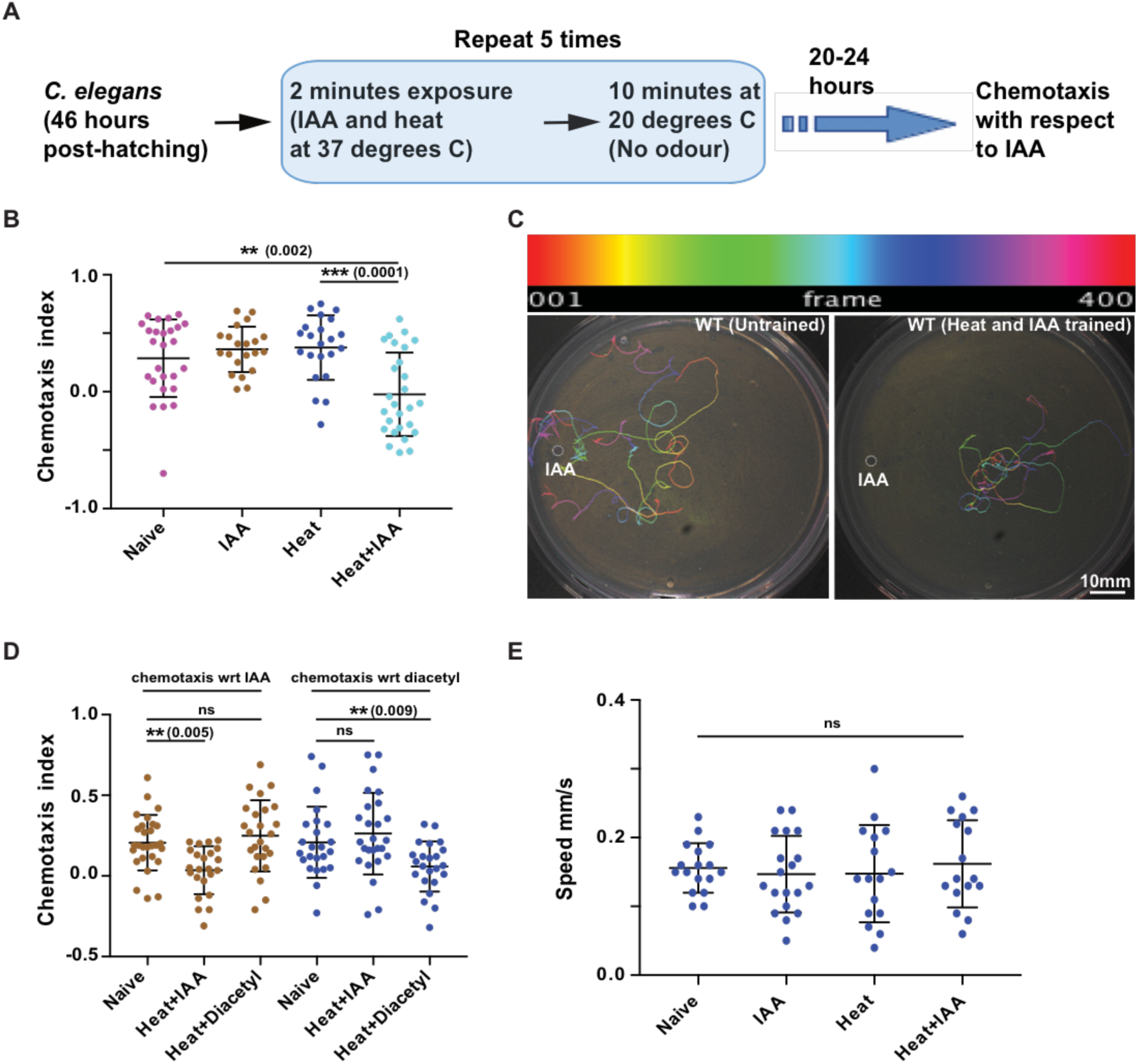
**(A)** Schematic flowchart of the ***C. elegans*** training routine followed in this study. **(B)** Scatter plot showing effect of IAA and heat on wild type (WT) worms**(C)** Representative images of typical chemotaxis tracks. The top panel shows the color-coding of tracks as a function of frame number. **(D)** Scatter plot indicating the specificity of learning. Animals were trained with Heat+IAA or Heat+Diacetyl and their chemotaxis indices were plotted. **(E)** Plot showing speed of worm movement when subjected to different training conditions. Error bars are SD. In graphs of all figures significant p values are added in brackets next to the significance stars (*).

#### Chemotaxis assay and behavioral analysis

Trained animals were kept at 20°C for 20-24 hours (h) unless mentioned otherwise. ***C. elegans*** were picked using an eyelash pick and allowed to crawl on an unseeded Nematode Growth Medium (NGM) plates for 30 seconds (s). Four to six animals were gently transferred to the center of 90mm NGM agar plates (without food). One µl of odorant (IAA (1%)/diacetyl (0.1%)) was kept at one end of the plate. Chemotaxis behavior was recorded for 10m using a 5 megapixel CMOS USB camera (Mightex) at 2 frames/s using a Mightex Camera Demo (v1.1.0) software. Recordings were done in a peltier cooled incubator at 20° Celsius. Video was analyzed using FIJI software (Schindelin et al., 2012). In order to quantify their chemotaxis behavior, a non-dimensional index based on individual ***C. elegans*** trajectory was used (Equation 1). For the attractant, displacement was positive if the animal traveled up the gradient while it was negative if the animal traveled down the gradient (Luo et al., 2014). To quantify the reversal behavior of the ***C. elegans*** along a gradient containing an attractive cue, IAA, a dimensionless index based on individual animal movement was used (Equation 2).

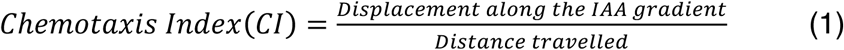

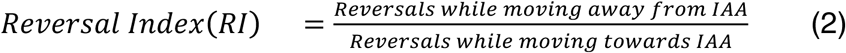

### 3.4 Quantitative PCR

Total RNA was isolated using Trizol from WT animals grown to young adult stage. 50ng of total RNA was used for cDNA synthesis. qPCR was performed by using a Qiagen SYBR real time PCR kit. Isoform specific primers were used for amplification of the different ***crh-1*** isoforms. The primers used are listed in in Table 1. qPCR was done using Roche Light Cycler 480. C_t_ values were calculated using equation 3 with the Roche software. The data was analyzed using ΔC method (Equation 4).

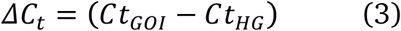

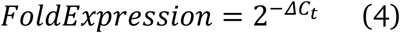

### 3.5 Fluorescence Microscopy

Young adult hermaphrodites were used for imaging. Animals were mounted on 2% agarose pads with 10mM sodium azide solution in M9 medium. Images were acquired on a Leica SP6 upright laser scanning confocal microscope using the 40X oil immersion objective lens. All images were processed and analyzed by FIJI.

### 3.6 Histamine supplementation

1M Histamine-dihydrochloride (Sigma-Aldrich, Cat No 53300) filter sterilized stock solution was prepared in distilled water. 10mM final working concentrations were used for all experiments. Histamine supplemented NGM plates were prepared as described previously (Pokala et al., 2014).

### 3.7 Optogenetics

The ***str-2*** promoter was used to drive AWC neuron-specific expression of Channelrhodopsin-2 (ChR2). Animals expressing ChR2 were grown on NGM agar plates seeded with OP50-containing 100μM all-trans retinal. The ***C. elegans*** were grown in the dark until late L4/early adult stages. The assay was performed on freshly seeded NGM plates. During the assay, ChR2 was excited by blue light (460-490 nm) using an epifluorescence unit (U-HGLGPS, Olympus) attached to the Nikon SMZ2000 microscope. Worms were illuminated with blue light for 3s and reversal events were quantified. If any worm executed reverse movement for one body length it was counted as a reversal event. Each worm was illuminated with blue light 8 times with 20s intervals between subsequent stimulation. Reversal probability was calculated by dividing number of reversal events with number of optogenetic stimulations. Each dot in the scatter plot indicates the reversal probability of a single worm under observation. The results were plotted as mean SD and evaluated using the standard Student’s t-test. The experimenter was blind to the genotypes of the strains while performing these experiments.

### 3.8 Calcium Imaging

The genetically encoded Ca^+2^ indicator GCaMP expressing strain ***prig-3***::GCaMP5 was used to visualize Ca^+2^ transients in the AVA command interneuron (Larsch et al., 2013). 0.2μl drop of polystyrene beads (0.1μm) was added on the top of 10% agarose pad (prepared in M9). A single worm was kept on the drop containing beads. The animals were immobilized by putting a circular cover glass over it. ChR2 was excited by blue light (460-490nm) using an epifluorescence unit (U-HGLGPS, Olympus) and simultaneously calcium transients were recorded for 3.0s at a speed of 10 frames/s. Images were captured using micro manager software (Edelstein et al. 2014). Image analysis was done using FIJI. A ROI was drawn over the AVA cell body. Fluorescence (F) was calculated by subtracting the background fluorescence (F_bkgd_) value from the mean fluorescence (F_mean_) of ROI. The fluorescence value was estimated for each frame by manual repositioning of the ROI. Calcium transients were plotted as ΔF/F_o_, where ΔF is the change in the fluorescence value (F) from its baseline fluorescence (F_o_). Fluorescence intensity of 2^nd^ frame (t=200ms) was taken as baseline fluorescence (F_o_).

### 3.9 CRISPR based genome editing

CRISPR was used to create ***crh-1*** isoform mutations as described previously (Dickinson and Goldstein, 2016). Selection Excision Cassette (SEC) from the plasmid pDD287 was cloned along-with flanking loxP sites into pPD95.75. The resulting plasmid was used to clone homology arms (500-1500bp) and the desired genetic modification using restriction enzyme based cloning methods. 20 base pair guide RNA was cloned into pRB1017 (Supplementary Figure 2). The plasmid mixture containing repair template (50ng/μl), sgRNA (20ng/μl), pJW1259 (50ng/μl), pCFJ90 (2.5ng/μl) and p***vha-6***::mCherry (10ng/μl) was injected into 20-30 adult hermaphrodite animals (containing 4-5 eggs). The Worms were kept at 20°C. After 60h of injection, the antibiotic hygromycin (250μg/ml) was added directly to the plates containing the injected worms. The ***C. elegans*** were left at 20°C for 10 days. After 10 days 10-15 non-fluorescent rollers were singled out onto regular NGM plates. To remove SEC 30-40 L1-L2 animals from plates with 100% roller progeny were kept at 34°C for 3-4h. Normal moving worms were isolated and target DNA was sequenced for desired modification (illustrated in Supplementary Figure 3).

### 3.10 Statistical analysis

Chemotaxis Index values of trained and untrained ***C. elegans*** were compared using the Student’s t-test. The ***crh-1*** isoform expression levels were compared using One way ANOVA. Error bars represent SD except for figure 5E (SEM). The level of significance was set as p<=0.05.

## 4. Results

### 4.1 Associative learning in *C. elegans* using isoamyl alcohol and high temperature

Previous studies have largely examined associative long-term memory (LTM) formation in ***C. elegans*** by pairing the presence/absence of food (unconditioned stimulus) with a variety of cues (Amano and Maruyama, 2011; Kauffman et al., 2011; Nishida et al., 2011). Under these previously reported training conditions ***C. elegans*** could retain memory for up to 40 hours (h) (Kauffman et al., 2011). Since studies have shown that starvation alone in the absence of any external stimulus is sufficient to induce the expression of CRH-1 in ***C. elegans*** (Suo et al., 2006, 2009), we were interested in developing training paradigm that was independent of the feeding state of the animal. To this end we developed a learning paradigm using isoamyl alcohol (IAA) and high temperature.

Worms were exposed to the vapors of IAA at high temperatures (37°C) for 2 minutes (m) followed by a rest period of 10m at 20°C, repeated five times (illustrated in Figure 1A). After 20-24h, the animals were allowed to crawl on a 90 mm NGM agar plate in response to IAA gradient. We observed significantly reduced chemotaxis index (CI) values for worms exposed to IAA at 37°C (Figures 1B, plate images shown in Figure 1C and Supplementary Figures 1A and B), while the CI values for worms exposed only to 37°C temperature or IAA alone were comparable to naïve animals (Figure 1B). To understand whether learning is specific to the cues provided during training, we trained worms with heat and IAA pairing while testing the LTM after 24h with chemotaxis w.r.t. diacetyl and vice versa. IAA when paired with heat resulted in reduced chemotaxis to IAA while chemotaxis to diacetyl was unaffected and vice versa (Figure 1D). To negate the potentially harmful effects of the chemicals/high temperature used in training we measured the average crawling speed of worms that were subjected to various training conditions, these animals crawled at speeds comparable to naïve animals (Figure 1E).

### 4.2 Long-term memory formation in *C. elegans* requires CRH-1c and CRH-1e

The ***creb1/crh-1*** gene is pleotropic in function, known to be involved in a variety of signaling pathways like cellular energy metabolism, ageing, memory formation, circadian rhythm etc. (Johannessen et al., 2004). Using multiple isoforms of a protein to achieve functional diversity is a widely used strategy during the course of molecular evolution (Graveley, 2001; Touriol et al., 2003; Perrin and Ervasti, 2010). In order to test if the above learning paradigm would allow us to parse out the isoform/s of CRH-1 required for long-term memory, we first tested the ***crh-1*** null mutant, ***crh-1(tz2) (***depicted in Figure 2A). We observed that trained wild type (WT) worms showed significantly reduced CI when compared to ***crh-1*** null mutants (Figure 2B).

**Figure 2.**
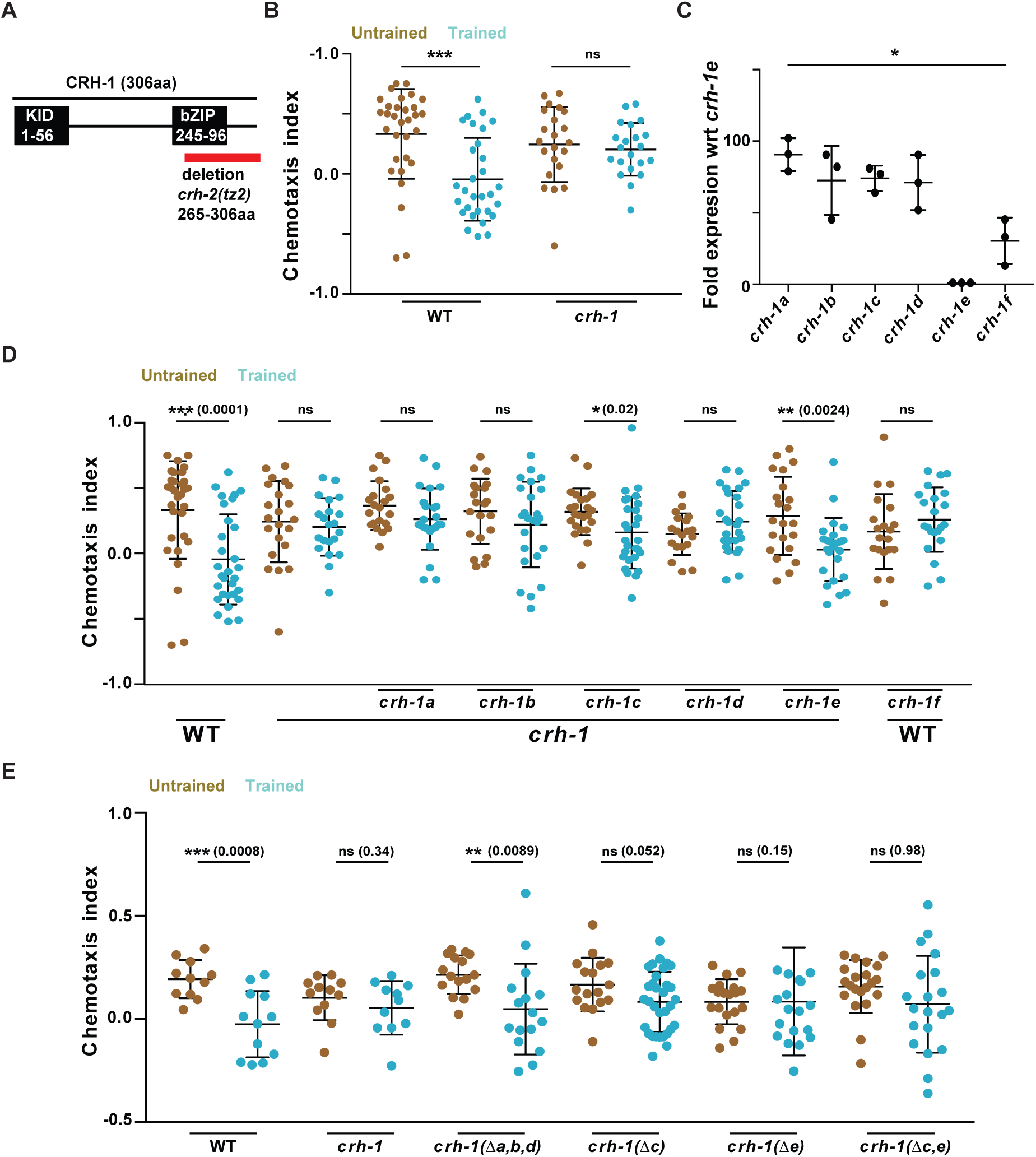
**(A)** Schematic showing ***C. elegans*** CRH-1e. Two major functional domains KID (kinase inducible domain) and bZIP (DNA binding domain) domains are depicted. The red bar shows the deletion found in the ***crh-1*** null mutant ***tz2***. This deletion affects all CRH-1 isoforms. **(B)** Scatter plot showing chemotaxis indices of WT and ***crh-1*** null mutants. **(C)** Graph showing quantitative PCR results comparing mRNA expression levels of different CRH-1 isoforms in young adult WT ***C. elegans***. **(D)** Scatter plot showing chemotaxis indices of WT, ***crh-1*** null mutants and pan neuronal CRH-1 rescue lines using different ***crh-1*** isoforms. **(E)** Scatter plot showing chemotaxis indices of WT, ***crh-1*** null mutants and ***crh-1*** isoform deletion lines. Error bars are SD.

The ***crh-1*** gene encodes seven different isoforms (***crh-1 a-g***) (Wormbase gene: WBGene00000793). We were able to clone cDNA for six of the seven isoforms (***crh-1 a-f***). We performed a quantitative PCR experiment to examine the expression levels of the different isoforms of CRH-1 during the early adult stage of worms. We observed strikingly variable expression levels for the various isoforms indicating a possibility of functional diversity among the different isoforms (Figure 2C). In order to further investigate the function of CRH-1 in memory, different CRH-1 isoforms i.e. ***crh-1 (a-f)*** were cloned under the pan-neuronal ***rab-3*** promoter and their ability to rescue the LTM defect in ***crh-1*** null mutants was tested (Figures 2D). Five of the six CRH-1 isoforms (CRH-1a-e) are full-length proteins having both the KID (Kinase Inducible Domain) and the bZIP (DNA binding) domain, these five isoforms differ only in the N-terminal 30 amino acids, while CRH-1f is a truncated protein lacking the N-terminal KID motif (Supplementary Figure 1C). We observed rescue of LTM formation in case of CRH-1c and CRH-1e expressing animals. Moreover, pan-neuronal expression of the repressor isoform CRH-1f in WT animals inhibited memory formation (Figure 2D). To negate the potential artifacts of spatiotemporal misexpression and/or over-expression due to extrachromosomal arrays, we generated CRISPR mutants for different ***crh-1*** isoforms and tested them for LTM defect (Supplementary Figures 2 and 3). Deletion of a, b and d isoforms by removing their first exon had no effect on LTM formation while worms without c and e isoforms were defective in LTM formation (Figure 2E).

### 4.3 Expression of CRH-1e in RIM or AVE neurons rescues the learning defects of *crh-1*mutants

Under stimulus deficient environments, the movement of ***C. elegans*** can be described as a random walk. The presence of an attractant/repellent introduces a bias in this random walk strategy whereby the animal can suppress or enhance its reversal and turn frequency depending on whether it is moving towards or away from the attractant/repellent (Pierce-Shimomura et al., 1999). In order to test if naïve ***crh-1*** null mutants showed any differences in their random walk pattern, we compared the reversal frequency of WT and ***crh-1*** null mutants while they were moving on an NGM agar plate in response to an IAA gradient. Our results indicated that ***crh-1*** null mutants were defective in modifying their reversal behavior in response to the varying attractant concentrations that they were experiencing while moving (Figure 3A). Reversal behavior in ***C. elegans*** is controlled by the command interneurons (AVA, AVD, AVE) and the RIM interneurons (Chalfie et al., 1985; Gray et al., 2005; Piggott et al., 2011). To test whether CRH-1e is required in these neurons we used a transgenic line expressing CRH-1e under the control of ***nmr-1*** promoter, which is expressed in five sets of neurons including AVA, AVD, AVE and RIM (Brockie et al., 2001). This line could completely rescue the reversal defect as well as the memory deficits observed in the ***creb1/crh-1*** null mutants (Figures 3A and B).

**Figure 3.**
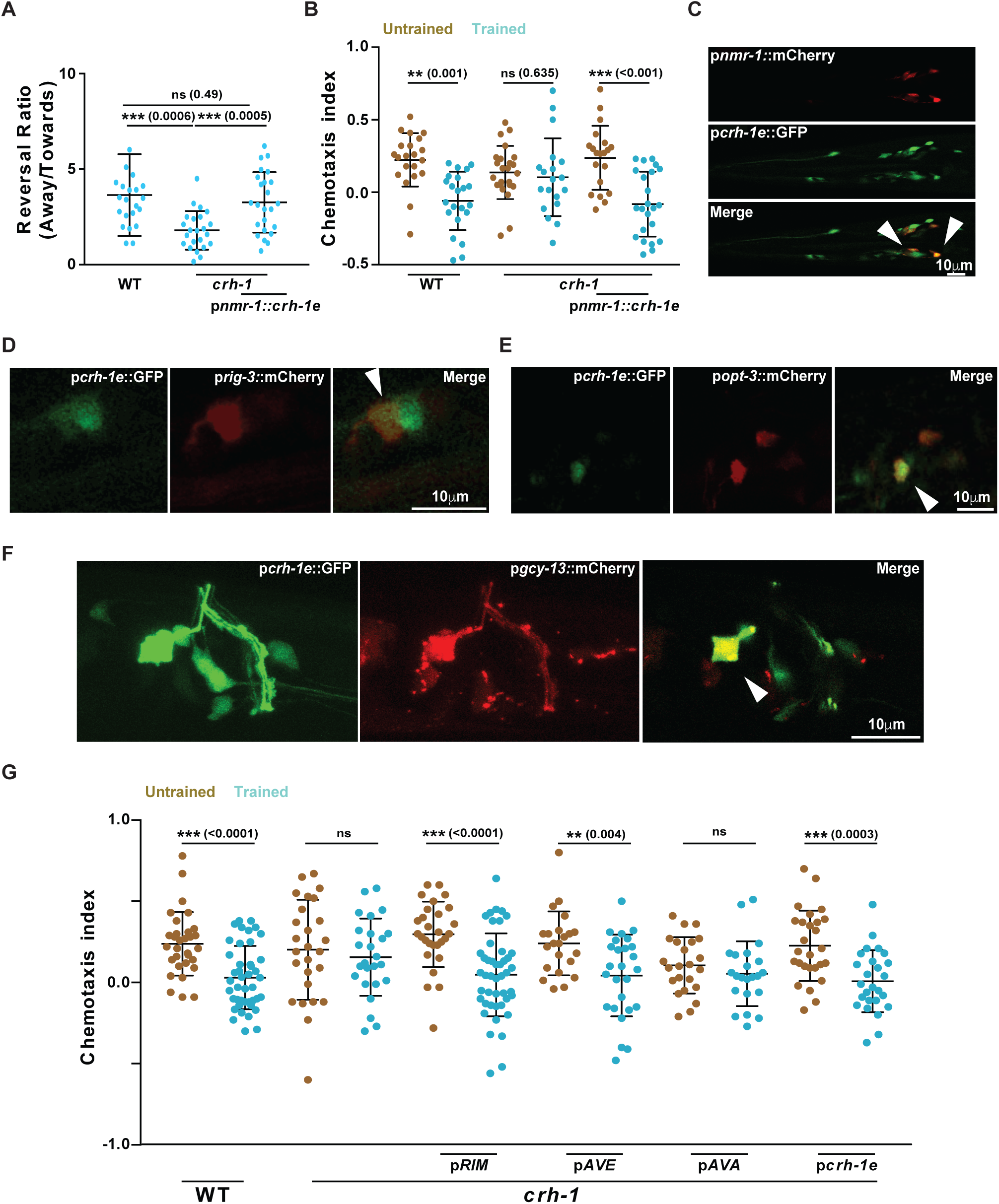
**(A)** Scatter plot showing reversal ratio of ***C. elegans*** while they were moving towards or away from the IAA point source. Each dot represents reversal events for individual animals counted for 10 minutes of chemotactic movement in response to IAA. **(B)** Chemotaxis indices for WT, ***crh-1*** and p***nmr-1::crh-1e*** rescue line. **(C)** Confocal microscope images showing overlapping expression of ***crh-1e*** promoter with neurons showing expression with the ***nmr-1*** promoter. Arrowheads indicate points of co-expression. **(D, E F)** Confocal microscope images showing overlapping expression of ***crh-1e*** promoter with neuron specific promoters; p***rig-3*** (AVA), p***opt-3*** (AVE) and p***gcy-13*** (RIM). **(G)** Scatter plot showing chemotaxis indices of WT, ***crh-1*** and rescue lines expressing CRH-1e under the promoters; ***gcy-13*** (RIM), ***opt-3*** (AVE), ***rig-3*** (AVA) and ***crh-1e***. Error bars are SD.

The presence of a unique 5’UTR on CRH-1e mRNA along with its low mRNA expression levels (Wormbase gene: WBGene00000793 and Figure 2C) suggests that its promoter sequence may be different from the other ***crh-1*** isoforms. We expressed GFP under the control of a 2.7 kb DNA sequence upstream of the CRH-1e translation start site (henceforth termed as p***crh-1e***). Consistent with remarkably low levels of expression in qPCR measurements (Figure 2C), we could observe the GFP expression in the p***crh-1e***::GFP line in only a few head neurons, expression in the head neurons was also seen previously by Kimura ***et al*** who had done in situ hybridization of the ***crh-1*** gene (Kimura et al., 2002). Colocalization experiments showed that the overlap in expression pattern of p***nmr-1*** was largely restricted to three pairs of neurons (Figure 3C). Based on the position of the overlapping neuron, we assessed that localization was seen in the AVA, AVE and RIM interneurons. The same was confirmed by co-expression experiments using neuron specific promoter marker lines for AVA (p***rig-3***::mCherry), AVE (p***opt-3***::mCherry) and RIM (p***gcy-13***::mCherry) (Fei et al., 2000; Ortiz et al., 2006; Feinberg et al., 2008). We found that ***pcrh-1e***::GFP localized to all these neurons (Figures 3D, E and F). To test where CRH-1e is required for memory formation we expressed CRH-1e under the control of ***crh-1e***, ***rig-3***, ***opt-3*** and ***gcy-13*** promoters. When tested for rescue of the associated memory phenotype with IAA and heat, we found that ***crh-1e, opt-3*** and the ***gcy-13*** promoters could significantly rescue the learning defects seen in ***crh-1*** mutants (Figure 3G). Expressing CRH-1e in AVA using the ***rig-3*** promoter did not rescue the associative memory defects of ***crh-1*** mutants (Figure 3G).

A recent study has shown that the CRH-1a isoform is broadly expressed along the ***C. elegans*** body and rescued the aging defects seen in ***crh-1*** mutants (Chen et al., 2016). We were interested in finding out if CRH-1a was expressed in the command interneurons and in the RIM interneuron. In order to look for the localization of the ***crh-1a*** promoter in these neurons we made a p***crh-1a***::GFP construct and look for the co-localization between p***crh-1a***::GFP and p***nmr-1***::mCherry. Our results indicated that there was no overlap between ***pcrh-1a*** expression and the command interneurons (data not shown) corroborating our hypothesis of functional diversity among CRH-1 isoforms.

### 4.4 CRH-1 is required at the time of training

To study the temporal requirement of CRH-1e, we asked whether the activity of the CRH-1e expressing neurons is required at the time of training. We expressed HisCl1 under the control of p***crh-1e*** in WT animals and silenced the HisCl1 expressing neurons by growing the worms on histamine containing plates while executing the training protocol (from 10m before training to 2h after training) or during chemotaxis (20h after training till the end of the assay). Memory formation was completely abolished in the animals with silenced neurons during training while silencing during chemotaxis resulted in defective chemotaxis even in naïve animals (Figure 4A).

**Figure 4.**
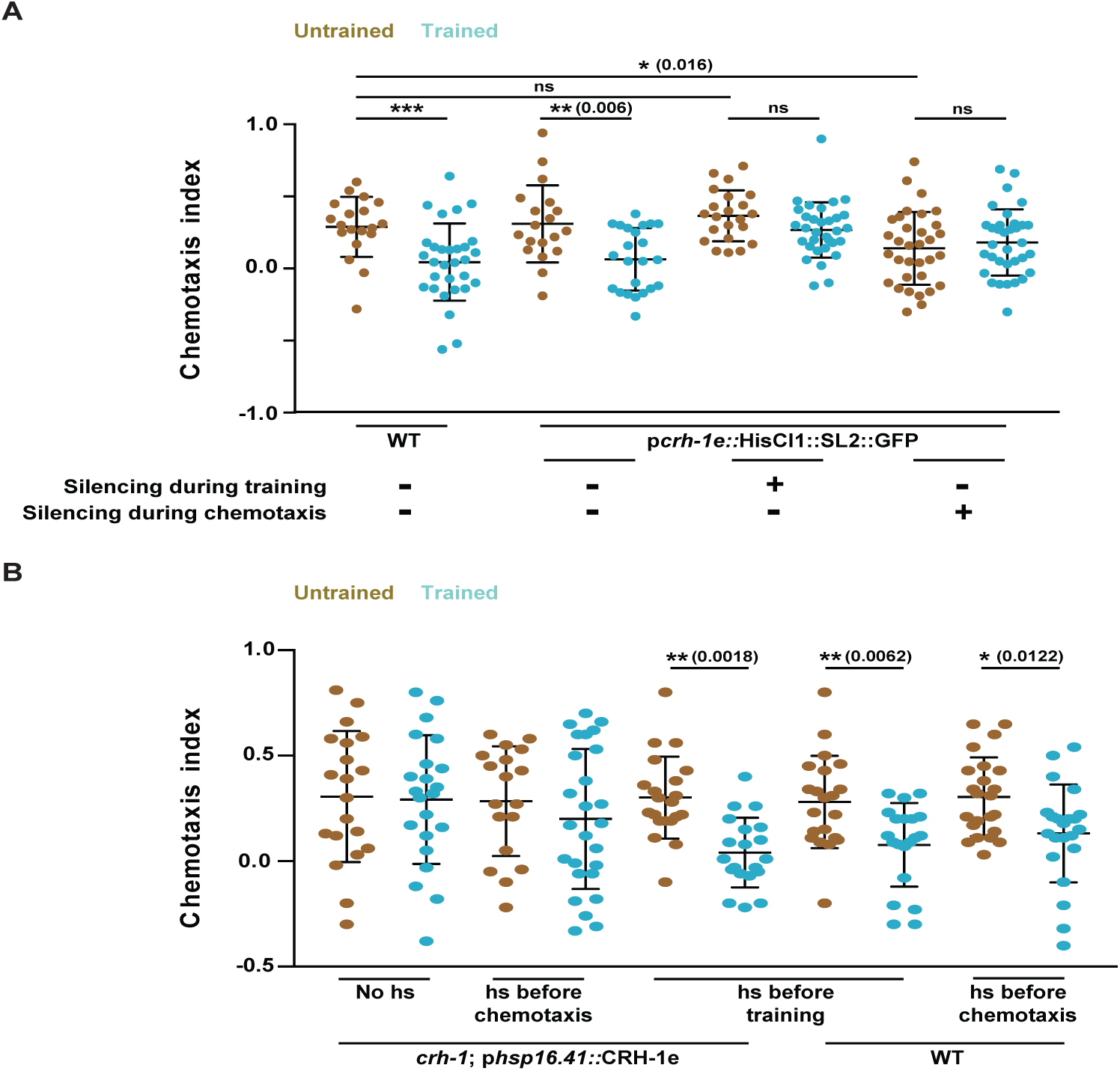
**(A)** Scatter plot showing chemotaxis indices of ***C. elegans*** under the condition of histamine mediated neuronal silencing**. (B)** Chemotaxis indices of WT and ***crh-1*** null mutants expressing CRH-1e under the control of the heat shock promoter ***hsp-16.41***. Error bars are SD.

To test the requirement of CRH-1e during the acquisition/consolidation phase of memory formation, we expressed CRH-1e under the control of the heat shock promoter (p***hsp16.41***) and used these animals to rescue the ***crh-1*** mutant phenotype. CRH-1e induction was done either 3h before training or 3h before chemotaxis. We observed that CRH-1e induction before training could largely rescue the learning defects, while induction before chemotaxis could not rescue the learning defect (Figure 4B). These results suggest that CRH-1e is required at the time of acquisition and/or consolidation phase of memory formation in *crh-1* mutant animals.

### 4.5 Lysine247 and lysine266 are key amino acid residues required for memory related functions of CRH-1

Most of the critical amino acid residues implicated in CREB1 activation are present in CRH-1a, b and d isoforms. CRH-1c and e differ from CRH-1a, b, d isoforms in having a much smaller and featureless N-terminal region. We hypothesized that the absence of these additional N-terminal residues is crucial for the functioning of CRH-1c and CRH-1e in LTM related processes. We added inert 3xHA tag to the N-terminal of CRH-1c and e (3xHA::CRH-1c,e) using CRISPR and tested these worms for the formation of LTM. Our experiments suggest that there was no significant difference in chemotaxis indices of trained and untrained 3xHA::CRH-1c,e animals (Figure 5A).

**Figure 5.**
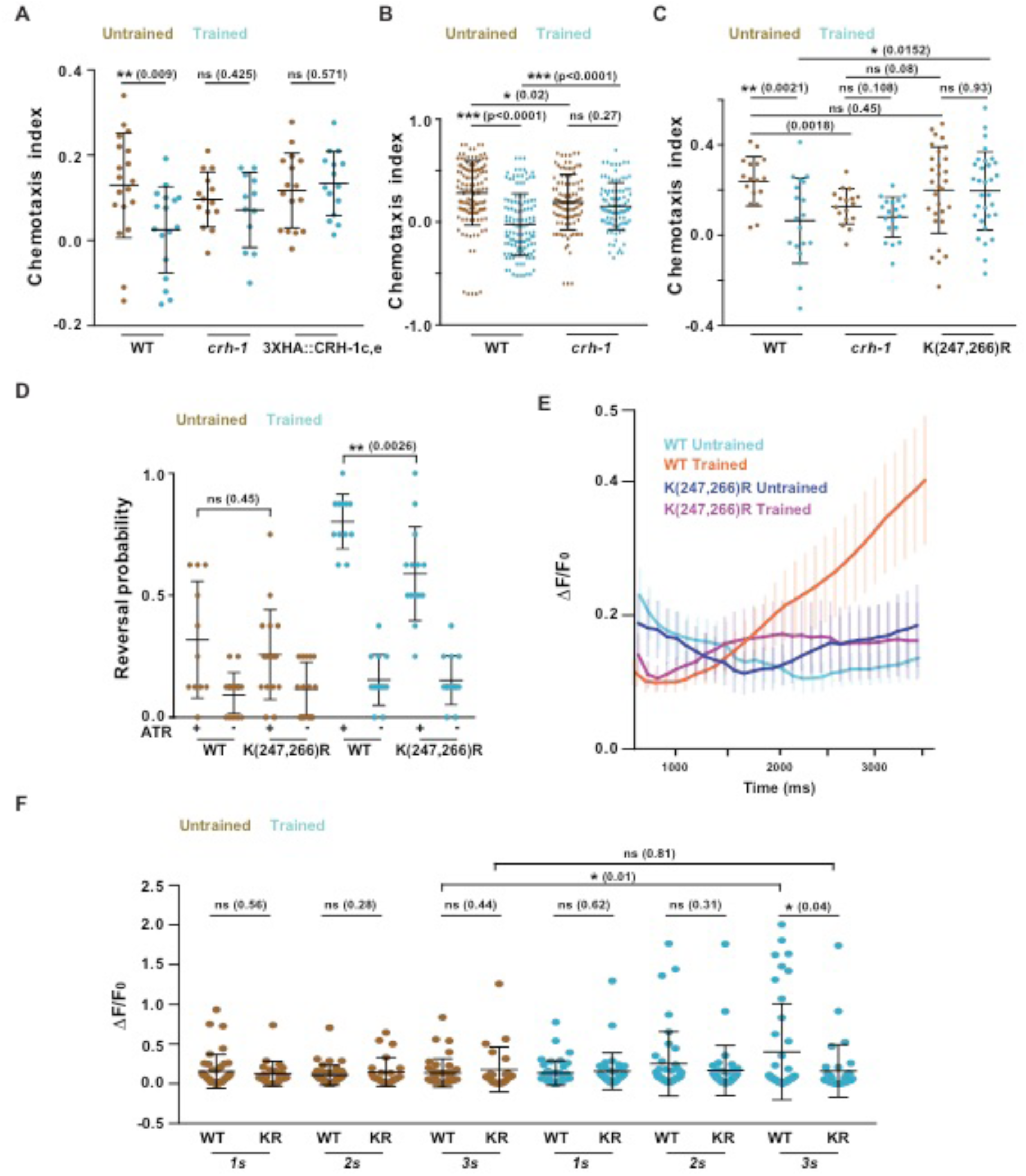
**(A)** Chemotaxis indices of WT, ***crh-1*** null mutants and N-terminal 3xHA tagged CRH-1c and CRH-1e. The tag was appended using CRISPR on N-terminal of CRH-1c and CRH-1e translation start codon. **(B)** Scatter plot showing collated chemotaxis index values of WT and ***crh-1*** null ***C. elegans*** used in this study. Data was pooled only form the experiments that used both strains. **(C)** Chemotaxis indices of WT, ***crh-1*** null mutants and ***crh-1***(K247R/K266R) mutants. **(D)** Reversal probability of ***C. elegans*** when AWC is excited with blue light. Blue light was illuminated for 3s and reversal events were quantified. If the worm executed reverse movement for one body length it was counted as a reversal event. Each ***C. elegans*** was illuminated with blue light 8 times with 20 second (s) intervals between subsequent stimulations. Each dot in the scatter plot indicates the reversal probability of a single animal under observation. **(E)** Plot showing Calcium traces of the AVA neuron in response to Channel Rhodopsin2 mediated AWC activation. Mean values are plotted against time. Error bars are SEM. N=25-41 for each genotype. **(F)** Scatter plot comparing AVA excitation in response to AWC activation at three time points. The Figure represents a subset of data (1s, 2s, 3s) from figure 5E. Error bars are SD except in figure 5E (SEM).

Null mutants of ***crh-1*** are defective in long-term memory formation. Naïve ***crh-1*** worms also show reduced chemotaxis indices compared to WT animals (Figure 5B). This defect is partially due to the inability of ***crh-1*** mutant animals to modulate reversal frequency in response to chemical gradients (Figure 3A). Therefore, observed learning defects in the ***crh-1*** mutant worms appear to be the compounded effect of learning deficit as well as defective native chemotactic behavior.

There are reports of dendritic localization of CREB1 (Crino et al., 1998). This increases the possibility of nuclear localization of CREB1 in response to dendritic stimulation. Moreover, SUMOylation dependent nuclear localization of CREB1 has been found to be critical for long-term memory formation in mouse (Chen et al., 2014). K285 and K304 have been identified as CREB1 SUMO acceptor sites in mouse (Comerford et al., 2003). To test for the possible involvement of these residues we generated K247R/K266R (corresponding to K285 and K304 in mouse CREB1) ***C. elegans*** using CRISPR. These mutant animals were trained and tested for the LTM phenotype. We observed that K247R/K266R mutant worms were defective in LTM formation while their naïve behavior was comparable to that seen in WT animals (Figure 5C).

The ability to modulate reversal frequency in response to chemical gradients is one of the primary factors mediating chemotactic behaviors in worms. We tested the effect of K247R/K266R mutations on the ability of ***C. elegans*** to initiate reversals in response to AWC activity. We optogenetically activated the AWC neuron and recorded the reversal probability of WT and K247R/K266R animals under naïve and trained conditions. Naïve WT and K247R/K266R animals showed similar reversal frequencies while trained worms showed significantly increased reversal frequencies with the K247R/K266R animals showing lower reversal probability in comparison to WT control animals (Figure 5D). To understand whether K247R/K266R mutations have any effect on AVA firing in response to AWC activity we optogenetically activated AWC for 3.0s while simultaneously imaging AVA using GCaMP-5G. While trained WT worms showed significant increase in the probability of AVA firing upon AWC activation there was no such increase seen in case of K247R/K266R (indicated as KR in Figure 5F) animals (Figure 5E and F). These data suggest the involvement of K247 and K266 residues in regulating the AVA firing and reversal probability of worms under specific training conditions.

## 5. Discussion

The role of CREB1 is well documented in the studies on learning and memory across phyla. It has been described as one of the main inducers of immediate early genes (IEGs) expressed in response to experience dependent neural activation (Alberini, 1999). However, functional pleiotropy of CREB1 remains a major bottleneck in studying CREB1 dependent IEGs that are activated specifically in response to experience dependent neural activation. Here, we describe the functional specialization of CRH-1 isoforms in ***C. elegans***. Our experiments show that ***C. elegans*** lacking CRH-1c and CRH-1e are defective in LTM formation while having normal native chemotaxis function. Null mutants of ***crh-1*** are defective in showing native as well as learned behavior. Restricted expression of CRH-1e in a small subset of neurons indicates functional specialization of the isoform in ***C. elegans.*** In ***Aplysia*** and ***Drosophila,*** it has been shown that different isoforms of CREB1 can repress and facilitate the process of memory formation (Yin et al., 1995; Yin and Tully, 1996; Bartsch et al., 1998). However, what separates CRH-1 isoforms from their CREB1 orthologs is their remarkable sequence similarity yet striking functional diversity. Spatial segregation is an efficient way of exploiting different functional properties of proteins. CRH-1e and CRH-1a have different expression patterns (this work and (Chen et al., 2016)) and this spatial segregation can explain their functional segregation. However, this might not be the whole story here since pan neuronal expression of CRH-1e cDNA could rescue the memory defects of ***crh-1*** null mutants; while expressing CRH-1a, the isoform that could rescue aging defect of null animals (Chen et al. 2016), could not rescue memory defects. Apart from the expression pattern differences CRH-1 isoforms also have small differences in their primary amino acid sequences, which are restricted to the N-terminal of the protein. It is possible that these small differences result in significantly altered tertiary structures as seen in other proteins (Goda et al., 2000; Korepanova et al., 2001), hence affecting activation dynamics or subcellular localization of the different protein isoforms (Crino et al., 1998). CRISPR mutants with K247R/K266R are defective in learned response while having normal native response to IAA gradient. SUMOylation of homologous residues has been shown to be associated with LTM defects in rats undergoing water maze test. It is yet to be seen whether K247 and K266 are targets for SUMOylation in ***C. elegans.***

Lau *et al* have shown that NMR-1 dependent associative memory defect is rescued by expressing NMR-1 in the RIM interneuron among other neurons (Lau et al. 2013). Previous work has shown that the glutamate receptor, GLR-1 function is necessary for LTM formation in ***C. elegans*** (Rose et al., 2003). AVE and RIM both express NMR-1 and GLR-1 and are connected to each other through gap junctions (Brockie et al., 2001; Kano et al., 2008; Kawano et al., 2011; Piggott et al., 2011; Lau et al., 2013). This along with our data makes it conceivable that RIM and AVE could be functioning together in the process of memory formation. Further, Xin ***et al*** have recently shown that the interneurons RIM and AIB are required for CRH-1 mediated imprinted memory formation, while the interneurons AIY and RIA are required for retrieval of the memory (Xin et al., 2016). Our results also show that presence of functional CRH-1e in RIM or AVE is sufficient for overcoming LTM formation defect in ***crh-1*** null worms. Both long-term associative memory and long-term habituation implicate CRH-1 in RIM neurons (Timbers and Rankin, 2011). Even though different neural circuits are activated by the training paradigms used in these studies, both paradigms operate in part by modulating the naïve reversal behavior in response to a cue. Consistent with these observations our experiments implicate K247 and K266 in modulating reversal probability in trained worms upon AWC activation through channelrhodopsin. Moreover, CRH-1 (K247R/K266R) trained animals have diminished AWC dependent AVA activation as measured by GCaMP5 mediated calcium imaging. This suggests that CRH-1 dependent genes might provide substrates for experience dependent modulation of AVA activity and reversal behaviors in ***C. elegans***.

AVA activity is highly correlated with reversal behavior in ***C. elegans*** (Gray et al., 2005; Ben Arous et al., 2010; Bhardwaj et al., 2018). The Ability to modulate AVA activity in response to sensory neuron activation provides for an important control point of behavioral modification through reversal modulation. In a parallel set of experiments, we have found that learning in worms is specific to the cue that was presented at the time of training. Therefore, given the strong correlation of AVA activity and reversal event it was surprising to find that expression of CRH-1e in AVE and RIM is sufficient to restore LTM defect in ***crh-1*** null mutants. AVA is heavily connected with AVE (44 incoming chemical synapses) and RIM (6 electrical synapses, 3 incoming chemical synapses) (White et al., 1986). Any change in AVE/RIM due to activity dependent CRH-1 activation in response to different sensory cues is likely to produce similar effect on AVA neurons. There are two probable explanations. One, the rescue of LTM defect that we observed in our experiments is an artifact of experimental methods used. Since specificity of promoters is largely determined by visible fluorescent protein reporter gene expression in a particular cell, it is likely that sub-visible expression at other sites is responsible for the LTM defect rescue. Second, learning but not its specificity is a function of CRH-1. If this is true, then it is likely that by studying CRH-1 dependent IEGs in ***C. elegans*** we will only be able to study somewhat mechanical aspect of memory manifesting at the terminal end of neural hierarchy just before behavior execution.

## Acknowledgments

YD and KB are Early Career and Intermediate Fellows of the DBT-Wellcome Trust India Alliance respectively and thank the Alliance for funding support (Grant nos. IA/E/13/1/501257 to YD and IA/I/12/1/500516 to KB). KB also acknowledges funding support from a DBT-IYBA grant (Grant no. BT/05/IYBA/2011) and from IISER Mohali. S.B was supported by KVPY fellowship, M.P and S.R. by DST-INSPIRE fellowships and S.T. by a CSIR-JRF. The *crh-1* mutant strains were provided by CGC, which is funded by National Institutes of Health Office of Research Infrastructure Program P40 OD010440. All the vectors used in this study were obtained from Addgene. The Indian Institute of Science Education and Research Mohali Confocal facility provided use of the confocal microscope. We thank members of the K.B. laboratory for comments on the manuscript; Cori Bargmann and Hernan Jaramillo for the p*rig-3*::GCCaMP5.0 array and strain CX15141 that we used to amplify HisCl1; Ankit Negi for routine help; Ashwani Bhardwaj and Saurabh Thapliyal for discussion, Arjit Kant Gupta for help in making chemotaxis video recording setup. We thank people contributing to and maintaining open source software; LibreOffice, Inkscape and Zotero. The authors declare no competing interests.

## 7. Supplementary figure

**Supplementary Figure 1.**
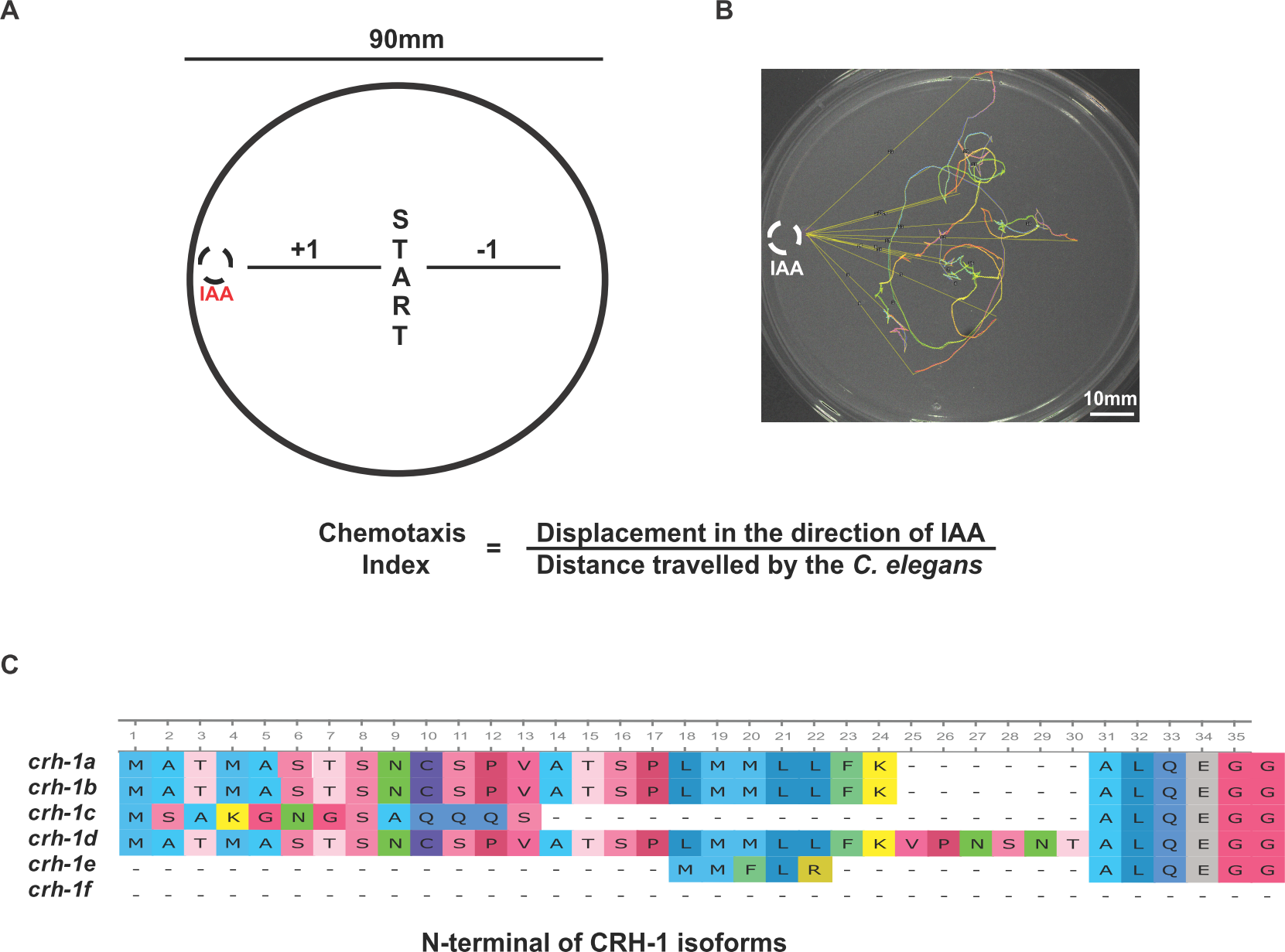
**(A)** Cartoon of a typical chemotaxis plate. 4-6 ***C. elegans*** are kept near the center of the plate and 1 µl of 1% IAA is kept at one end of the plate (dashed circle). A video is recorded as the animals move for 10 minutes using a 5 megapixel USB camera at 2 fps. ***C. elegans*** tracks are analyzed and the chemotaxis index is calculated. **(B)**Cartoon of calculation of ***C. elegans*** displacement using worm tracks. The worm tracks are generated by FIJI using temporal color code. The distance travelled by the worm is measured by following the track using the segmented line tool in FIJI. In order to measure the displacement in the direction of the attractant, the shortest distance from the end of the track to the source of the attractant is measured and this value is subtracted from the shortest distance from the start of the track to the source of the attractant. The chemotaxis index is calculated by dividing the displacement in the direction of the attractant by the total distance travelled by the ***C. elegans***. **(C)** This panel indicates sequence alignments of CRH-1 isoforms a-f using MUSCLE. The N-terminal part of the protein having maximum sequence variation is shown. The alignment was generated using UGENE software.

**Supplementary Figure 2:**
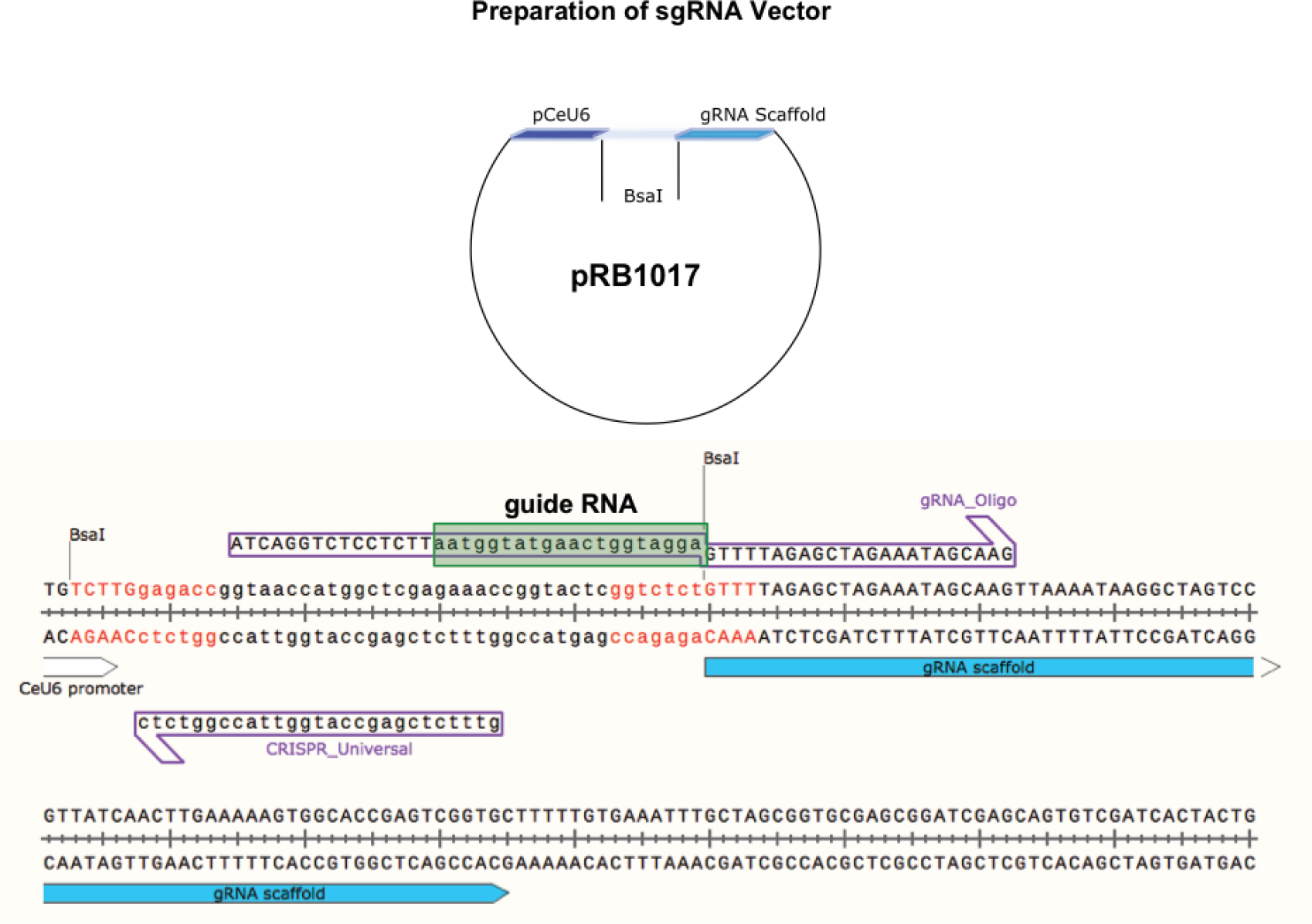
The vector used and sequence indicating the guide RNA for CRISPR mediated sequence changes implemented in this study.

**Supplementary Figure 3:**
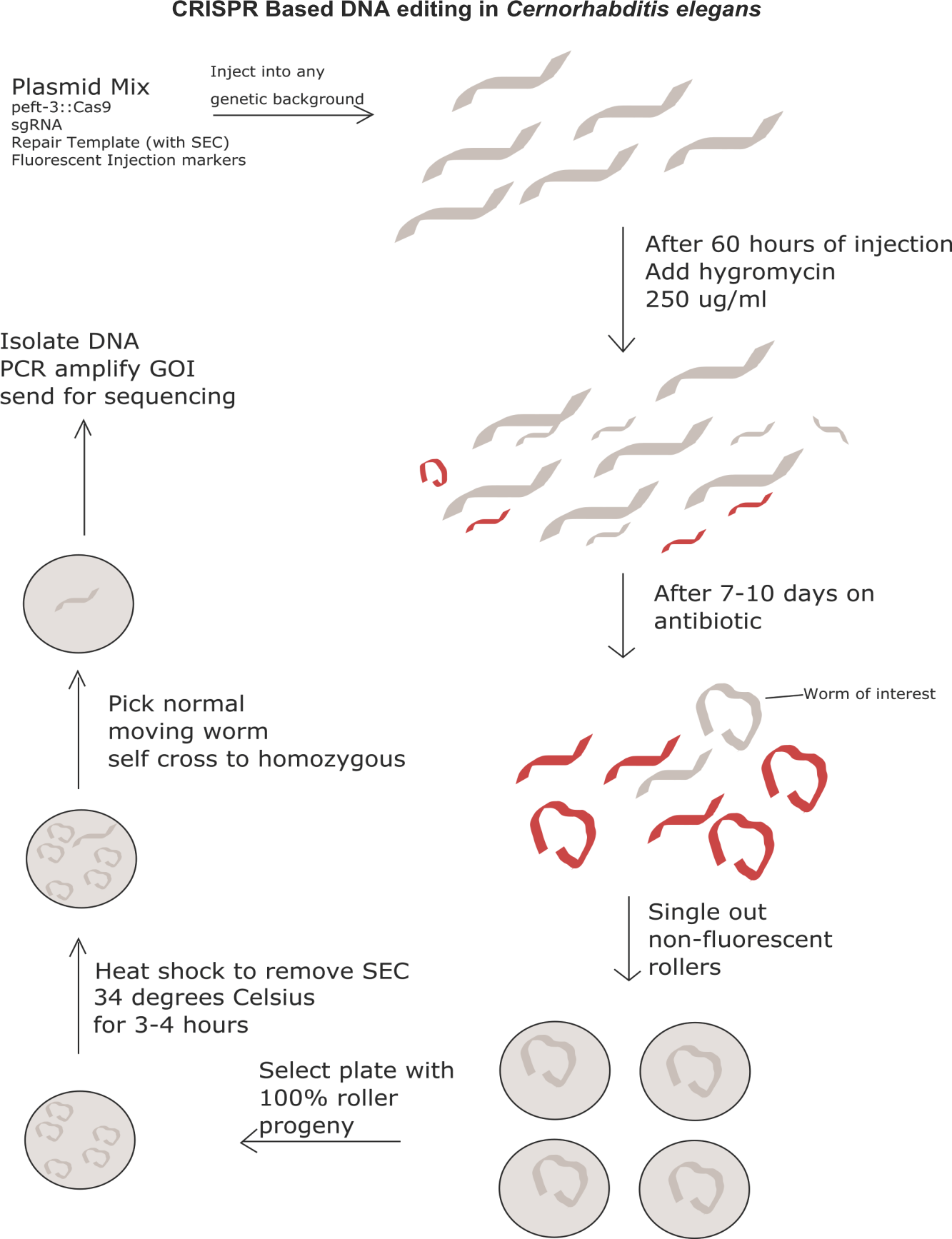
A schematic indicating how CRISPR mediated gene editing was done in this study.

Author contributions
YD designed and performed experiments, analyzed data, andwrote the manuscript; SR, ST, SB and MP performed experiments and analyzed data; KB supervisedand helped with editing the manuscript

## Bibliography

Alberini CM (1999) Genes to remember. J Exp Biol 202:2887–2891.

Amano H, Maruyama IN (2011) Aversive olfactory learning and associative long-term memory in Caenorhabditis elegans. Learn Mem 18:654–665.

Ardiel EL, Rankin CH (2010) An elegant mind: Learning and memory in Caenorhabditis elegans. Learn Mem 17:191–201.

Bargmann CI (2006) Chemosensation in C. elegans. In: Wormbook Available at: http://www.ncbi.nlm.nih.gov/books/NBK19746/ [Accessed January 22, 2015].

Bartsch D, Casadio A, Karl KA, Serodio P, Kandel ER (1998) CREB1 Encodes a Nuclear Activator, a Repressor, and a Cytoplasmic Modulator that Form a Regulatory Unit Critical for Long-Term Facilitation. Cell 95:211–223.

Bateson P, Mameli M (2007) The innate and the acquired: Useful clusters or a residual distinction from folk biology? Dev Psychobiol 49:818–831.

Ben Arous J, Tanizawa Y, Rabinowitch I, Chatenay D, Schafer WR (2010) Automated imaging of neuronal activity in freely behaving Caenorhabditis elegans. J Neurosci Methods 187:229–234.

Bhardwaj A, Thapliyal S, Dahiya Y, Babu K (2018) FLP-18 functions through the G-protein coupled receptors NPR-1 and NPR-4 to modulate reversal length in Caenorhabditis elegans. J Neurosci:1955–17.

Brenner S (1974) The genetics of Caenorhabditis elegans. Genetics 77:71–94.

Brockie PJ, Mellem JE, Hills T, Madsen DM, Maricq AV (2001) The C. elegans glutamate receptor subunit NMR-1 is required for slow NMDA-activated currents that regulate reversal frequency during locomotion. Neuron 31:617–630.

Chalfie M, Sulston JE, White JG, Southgate E, Thomson JN, Brenner S (1985) The neural circuit for touch sensitivity in Caenorhabditis elegans. J Neurosci 5:956–964.

Chen Y-C, Chen H-J, Tseng W-C, Hsu J-M, Huang T-T, Chen C-H, Pan C-L (2016) A C. elegans Thermosensory Circuit Regulates Longevity through crh-1/CREB-Dependent flp-6 Neuropeptide Signaling. Dev Cell Available at: http://linkinghub.elsevier.com/retrieve/pii/S1534580716305962 [Accessed October 11, 2016].

Chen Y-C, Hsu W-L, Ma Y-L, Tai DJC, Lee EHY (2014) CREB SUMOylation by the E3 Ligase PIAS1 Enhances Spatial Memory. J Neurosci 34:9574–9589.

Comerford KM, Leonard MO, Karhausen J, Carey R, Colgan SP, Taylor CT (2003) Small ubiquitin-related modifier-1 modification mediates resolution of CREB-dependent responses to hypoxia. Proc Natl Acad Sci 100:986–991.

Crino P, Khodakhah K, Becker K, Ginsberg S, Hemby S, Eberwine J (1998) Presence and phosphorylation of transcription factors in developing dendrites. Proc Natl Acad Sci 95:2313–2318.

Dash PK, Hochner B, Kandel ER (1990) Injection of the cAMP-responsive element into the nucleus of Aplysia sensory neurons blocks long-term facilitation. Nature 345:718–721.

Dickinson DJ, Goldstein B (2016) CRISPR-Based Methods for Caenorhabditis elegans Genome Engineering. Genetics 202:885–901.

Fei YJ, Romero MF, Krause M, Liu JC, Huang W, Ganapathy V, Leibach FH (2000) A novel H(+)-coupled oligopeptide transporter (OPT3) from Caenorhabditis elegans with a predominant function as a H(+) channel and an exclusive expression in neurons. J Biol Chem 275:9563–9571.

Feinberg EH, VanHoven MK, Bendesky A, Wang G, Fetter RD, Shen K, Bargmann CI (2008) GFP Reconstitution Across Synaptic Partners (GRASP) Defines Cell Contacts and Synapses in Living Nervous Systems. Neuron 57:353–363.

Flavell SW, Greenberg ME (2008) Signaling mechanisms linking neuronal activity to gene expression and plasticity of the nervous system. Annu Rev Neurosci 31:563–590.

Goda S, Takano K, Yamagata Y, Katakura Y, Yutani K (2000) Effect of extra N-terminal residues on the stability and folding of human lysozyme expressed in Pichia pastoris. Protein Eng 13:299–307.

Graveley BR (2001) Alternative splicing: increasing diversity in the proteomic world. Trends Genet 17:100–107.

Gray JM, Hill JJ, Bargmann CI (2005) A circuit for navigation in Caenorhabditis elegans. Proc Natl Acad Sci U S A 102:3184–3191.

Hart A (2006) Behavior. WormBook Available at: http://www.wormbook.org/chapters/www_behavior/behavior.html [Accessed January 23, 2015].

Iino Y, Yoshida K (2009) Parallel Use of Two Behavioral Mechanisms for Chemotaxis in Caenorhabditis elegans. J Neurosci 29:5370–5380.

Johannessen M, Delghandi MP, Moens U (2004) What turns CREB on? Cell Signal 16:1211– 1227.

Kano T, Brockie PJ, Sassa T, Fujimoto H, Kawahara Y, Iino Y, Mellem JE, Madsen DM, Hosono R, Maricq AV (2008) Memory in Caenorhabditis elegans Is Mediated by NMDA-Type Ionotropic Glutamate Receptors. Curr Biol 18:1010–1015.

Kauffman A, Parsons L, Stein G, Wills A, Kaletsky R, Murphy C (2011) C. elegans positive butanone learning, short-term, and long-term associative memory assays. J Vis Exp JoVE.

Kauffman AL, Ashraf JM, Corces-Zimmerman MR, Landis JN, Murphy CT (2010) Insulin signaling and dietary restriction differentially influence the decline of learning and memory with age. PLoS Biol 8:e1000372.

Kawano T, Po MD, Gao S, Leung G, Ryu WS, Zhen M (2011) An imbalancing act: gap junctions reduce the backward motor circuit activity to bias C. elegans for forward locomotion. Neuron 72:572–586.

Kimura Y, Corcoran EE, Eto K, Gengyo-Ando K, Muramatsu M, Kobayashi R, Freedman JH, Mitani S, Hagiwara M, Means AR, others (2002) A CaMK cascade activates CRE-mediated transcription in neurons of Caenorhabditis elegans. EMBO Rep 3:962–966.

Korepanova A, Douglas C, Leyngold I, Logan TM (2001) N0terminal extension changes the folding mechanism of the FK5060binding protein. Protein Sci 10:1905–1910.

Larsch J, Flavell SW, Liu Q, Gordus A, Albrecht DR, Bargmann CI (2015) A Circuit for Gradient Climbing in C. elegans Chemotaxis. Cell Rep 12:1748–1760.

Larsch J, Ventimiglia D, Bargmann CI, Albrecht DR (2013) High-throughput imaging of neuronal activity in Caenorhabditis elegans. Proc Natl Acad Sci 110:E4266–E4273.

Lau HL, Timbers TA, Mahmoud R, Rankin CH (2013) Genetic dissection of memory for associative and non-associative learning in Caenorhabditis elegans. Genes Brain Behav 12:210–223.

Luo L, Wen Q, Ren J, Hendricks M, Gershow M, Qin Y, Greenwood J, Soucy ER, Klein M, Smith-Parker HK, Calvo AC, Colón-Ramos DA, Samuel ADT, Zhang Y (2014) Dynamic Encoding of Perception, Memory, and Movement in a C. elegans Chemotaxis Circuit. Neuron 82:1115–1128.

Mello CC, Kramer JM, Stinchcomb D, Ambros V (1991) Efficient gene transfer in C.elegans: extrachromosomal maintenance and integration of transforming sequences. EMBO J 10:3959–3970.

Nishida Y, Sugi T, Nonomura M, Mori I (2011) Identification of the AFD neuron as the site of action of the CREB protein in Caenorhabditis elegans thermotaxis. EMBO Rep 12:855– 862.

Ortiz CO, Etchberger JF, Posy SL, Frøkjaer-Jensen C, Lockery S, Honig B, Hobert O (2006) Searching for neuronal left/right asymmetry: genomewide analysis of nematode receptor-type guanylyl cyclases. Genetics 173:131–149.

Pereira S, Kooy D van der (2012) Two Forms of Learning following Training to a Single Odorant in Caenorhabditis elegans AWC Neurons. J Neurosci 32:9035–9044.

Perrin BJ, Ervasti JM (2010) The actin gene family: Function follows isoform. Cytoskeleton 67:630–634.

Pierce-Shimomura JT, Morse TM, Lockery SR (1999) The fundamental role of pirouettes in Caenorhabditis elegans chemotaxis. J Neurosci 19:9557–9569.

Piggott BJ, Liu J, Feng Z, Wescott SA, Xu XZS (2011) The Neural Circuits and Synaptic Mechanisms Underlying Motor Initiation in C. elegans. Cell 147:922–933.

Pokala N, Liu Q, Gordus A, Bargmann CI (2014) Inducible and titratable silencing of Caenorhabditis elegans neurons in vivo with histamine-gated chloride channels. Proc Natl Acad Sci 111:2770–2775.

Rose JK, Kaun KR, Chen SH, Rankin CH (2003) GLR-1, a non-NMDA glutamate receptor homolog, is critical for long-term memory in Caenorhabditis elegans. J Neurosci 23:9595– 9599.

Schindelin J, Arganda-Carreras I, Frise E, Kaynig V, Longair M, Pietzsch T, Preibisch S, Rueden C, Saalfeld S, Schmid B, Tinevez J-Y, White DJ, Hartenstein V, Eliceiri K, Tomancak P, Cardona A (2012) Fiji: an open-source platform for biological-image analysis. Nat Methods 9:676–682.

Silva AJ, Kogan JH, Frankland PW, Kida S (1998) CREB and memory. Annu Rev Neurosci 21:127–148.

Suo S, Culotti JG, Tol HHMV (2009) Dopamine counteracts octopamine signalling in a neural circuit mediating food response in C. elegans. EMBO J 28:2437–2448.

Suo S, Kimura Y, Van Tol HHM (2006) Starvation induces cAMP response element-binding protein-dependent gene expression through octopamine-Gq signaling in Caenorhabditis elegans. J Neurosci Off J Soc Neurosci 26:10082–10090.

Tierney AJ (1986) The evolution of learned and innate behavior: Contributions from genetics and neurobiology to a theory of behavioral evolution. Anim Learn Behav 14:339– 348.

Timbers TA, Rankin CH (2011) Tap withdrawal circuit interneurons require CREB for long-term habituation in Caenorhabditis elegans. Behav Neurosci 125:560–566.

Touriol C, Bornes S, Bonnal S, Audigier S, Prats H, Prats A-C, Vagner S (2003) Generation of protein isoform diversity by alternative initiation of translation at non-AUG codons. Biol Cell 95:169–178.

White JG, Southgate E, Thomson JN, Brenner S (1986) The structure of the nervous system of the nematode Caenorhabditis elegans. Philos Trans R Soc Lond B Biol Sci 314:1–340.

Xin J, Pokala N, Bargmann CI (2016) Distinct Circuits for the Formation and Retrieval of an Imprinted Olfactory Memory. Cell 164:632–643.

Yin JC, Del Vecchio M, Zhou H, Tully T (1995) CREB as a memory modulator: induced expression of a dCREB2 activator isoform enhances long-term memory in Drosophila. Cell 81:107–115.

Yin JC, Tully T (1996) CREB and the formation of long-term memory. Curr Opin Neurobiol 6:264–268.

